# Phylogeographic structure of the dunes sagebrush lizard, an endemic habitat specialist

**DOI:** 10.1101/2020.06.23.168088

**Authors:** Lauren M. Chan, Charles W. Painter, Michael T. Hill, Toby J. Hibbitts, Daniel J. Leavitt, Wade A. Ryberg, Danielle Walkup, Lee A. Fitzgerald

**Author notes:** deceased (Charles W. Painter).

## Abstract

Phylogeographic divergence and population genetic diversity within species reflect the impacts of habitat connectivity, demographics, and landscape level processes in both the recent and distant past. Characterizing patterns of differentiation across the geographic range of a species provides insight on the roles of organismal and environmental traits, on evolutionary divergence, and future population persistence. This is particularly true of habitat specialists where habitat availability and resource dependence may result in pronounced genetic structure as well as increased population vulnerability. We use DNA sequence data as well as microsatellite genotypes to estimate range-wide phylogeographic divergence, historical population connectivity, and historical demographics in an endemic habitat specialist, the dunes sagebrush lizard (*Sceloporus arenicolus*). This species is found exclusively in dune blowouts and patches of open sand within the shinnery oak-sand dune ecosystem of southeastern New Mexico and adjacent Texas. We find evidence of phylogeographic structure consistent with breaks and constrictions in suitable habitat at the range-wide scale. In addition, we find support for a dynamic and variable evolutionary history across the range of *S. arenicolus*. Populations in the Monahans Sandhills have deeply divergent lineages consistent with long-term demographic stability. In contrast, populations in the Mescalero Sands are not highly differentiated, though we do find evidence of demographic expansion in some regions and relative demographic stability in others. Phylogeographic history and population genetic differentiation in this species has been shaped by the configuration of habitat patches within a geologically complex and historically dynamic landscape. Our findings identify regions as genetically distinctive conservation units as well as underscore the genetic and demographic history of different lineages of *S. arenicolus*.

## Introduction

Patterns of population genetic diversity within species are shaped by both evolutionary and contemporary history (Rissler, 2016). Though anthropogenic changes to landscapes alter patterns of connectivity that can result in the divergence or coalescence of populations, these processes take place on a background of evolutionary history determined by chance, species’ life history, and also geologic and climatic changes. Characterizing this evolutionary history, and identifying the role that organismal traits, evolutionary processes, and ecological conditions have on patterns of phylogeographic divergence adds to our understanding of evolution, and is also fundamental to conserving evolutionary potential in the face of anthropogenic disturbance and climate change (Olivieri et al., 2015).

The phylogeographic history of species can reflect the roles that habitat connectivity, gene flow, and population stability have played in a species’ evolutionary persistence. Some species may be characterized by deeply divergent lineages, suggesting a history of limited dispersal and low connectivity among sites (e.g., Richmond et al., 2013, 2014; Chan et al., 2013), especially in ecosystems with steep environmental gradients and discontinuous habitat (Vandergast et al., 2008). Plant and animal taxa in naturally fragmented landscapes, for example, can exhibit strong patterns of genetic population structure with selection favoring limited dispersal. Phylogeographic analyses of *Stenopelmatus* species (Jerusalem crickets) in southwestern North America, for example, revealed limited dispersal among populations, and identified a recent response to anthropogenic change (Vandergast et al., 2007). A meta-analysis of genetic diversity among 21 species of terrestrial animals identified hotspots of genetic diversity that may also be regions with high levels of trait divergence due to natural selection (Vandergast et al., 2008). Alternatively, populations may be only weakly divergent across a species’ range indicating high connectivity (e.g. Lippé et al., 2006; Chan and Zamudio, 2009) even in the face of strong local dynamics (e.g. Pierson et al., 2013). Identifying evolutionary scenarios and processes that have resulted in particular phylogeographic patterns can help us disentangle processes that underlie population genetic divergence from those that maintain genetic diversity. Understanding the drivers of population genetic structure across the range of a species can also help us predict the response to loss of habitat and the overall vulnerability of species to anthropogenic landscape change.

Ecological specialists can have greater population genetic and phylogeographic structure than generalists because individuals and populations may be restricted to spatially isolated patches of suitable habitat. (Roderick et al., 2012; Schär et al., 2018; Wort et al., 2019). Ecological specialists may have narrow physiological tolerances, specific habitat requirements, and be locally abundant but rare at regional scales (Devictor et al., 2008). Habitat specialists use specific landscape features and vegetation associations within their range, and often possess eco-morphological and behavioral adaptations (Miles, 1994a, 1994b). Traits that make habitat specialists well-suited for a narrow habitat niche also tend to make them relatively poor dispersers (Clobert et al., 2012). Low tolerance for unsuitable landscapes is expected to restrict movements among isolated patches of preferred habitat. Ecological studies focusing on the demography and distribution of habitat specialists have found they are sensitive to landscape fragmentation (Leavitt and Fitzgerald, 2013; Walkup et al., 2017).

Local processes are often linked to patterns observed across long-term, evolutionary time scales and at broader spatial scales (see reviews by Cutter, 2013; Rissler, 2016). Thus, in specialists with strict habitat specificity and limited dispersal among populations, we might expect phylogeographic structure to reflect historical patterns of divergence and low population connectivity overall (e.g., Roderick et al., 2012). Alternatively, habitat specialists may have well-connected populations throughout their range, indicating a strong role for dispersal and migration that counters the divergence of potentially isolated local populations across longer time-scales (Pierson et al., 2013). Characterizing evolutionary patterns of divergence and historical demographics in habitat specialists can help us predict the role that short and long-term dynamics play in shaping population genetic structure. In addition, describing spatial patterns of diversity and identifying independent evolutionary units, historical barriers to gene flow, bottlenecks and founder events, and regions of high connectivity allows the effects of contemporary pressures to be disentangled from historical drivers and also provides important information for the future management and conservation of species.

The dunes sagebrush lizard, *Sceloporus arenicolus*, is endemic to the Mescalero and Monahans Sandhills ecosystem of southeastern New Mexico and adjacent Texas (Fitzgerald and Painter, 2009; Laurencio and Fitzgerald, 2010). This species is part of the *Sceloporus graciosus* clade (Chan et al., 2013), but in contrast to other members of this group which tend to be geographically widespread generalists, *S. arenicolus* is a habitat specialist. Within this ecosystem, it only uses shinnery-oak sand dune formations with interconnected dune blowouts (sandy depressions created by wind) and in some cases shinnery hummocks in dunes with steep slopes (Fitzgerald et al., 1997; Laurencio and Fitzgerald, 2010; Hibbitts et al., 2013). In the Mescalero-Monahans Sandhills Ecosystem, dune blowouts are emergent landforms that are maintained by the interactions among wind, moving sand, and the shinnery oak (*Quercus havardii*) which stabilizes the dunes (Ryberg and Fitzgerald, 2016). Individual *S. arenicolus* lizards demonstrate a nested hierarchy of habitat selection (Fitzgerald et al., 1997), selecting for thermally suitable microhabitats and having preference for relatively large dune blowouts. A sand-diving species, they do not occur in areas with relatively fine sand (Fitzgerald et al. 1997; Ryberg and Fitzgerald 2015). At the highest level of habitat selection, they are endemic to the narrowly distributed Mescalero-Monahans Sandhills (Fitzgerald and Painter, 2009).

Specialists can reach high population densities in their preferred habitat, and can outcompete generalists in the same area even in some degraded habitats (Brown, 1984; Attum et al., 2006). This is true too for *S. arenicolus*, where populations of this ecological specialist thrive where the configuration of key landscape features supports larger groups of interacting individuals, defined as neighborhoods (*sensu* Wright, 1946; Ryberg et al., 2013). Diffusion dispersal throughout interconnected areas of suitable habitat appear key to maintaining populations in contiguous habitat over the long term (Ryberg et al., 2013). The quantity of habitat is positively correlated with the quality of habitat (Smolensky and Fitzgerald, 2011), and the occurrence of *S. arenicolus* is associated with relatively large core areas of shinnery oak dunes.

Since at least the 1930s, anthropogenic disturbances from herbicide spraying, oil and gas mining, and more recently, sand-mining, have resulted in fragmentation and degradation of the shinnery oak dunes. Long-term monitoring and extensive fieldwork also demonstrate that fragmentation of the shinnery oak dunelands leads directly to population collapse because quality of habitat tends to degrade in response to fragmentation, and dispersal is disrupted (Leavitt and Fitzgerald, 2013; Walkup et al., 2017).

To adequately inform conservation and management actions, it is necessary to understand the evolutionary history of this species at both broad and fine-scales throughout the range. Previous genetic work confirmed that at broad spatial scales, *S. arenicolus* is comprised of at least three distinct genetic groups (Chan et al., 2009). However, it is unclear where these genetic breaks occur geographically and whether they coincide with putative natural or man-made barriers to movement. The purpose of this study is to characterize the evolutionary history of the dunes sagebrush lizard using complete geographic and genetic sampling. To identify evolutionary distinct geographic lineages and to reconstruct the population history of these lineages, we evaluate mitochondrial and nuclear sequence data as well as multilocus microsatellite genotypes. Sampling for individuals occurred evenly throughout the entire known range of this endemic and threatened lizard.

## Materials and Methods

### Sampling

We surveyed for *Sceloporus arenicolus* throughout their range (Figure 1). Liver or muscle tissue was collected from vouchered specimens deposited in the Biodiversity Research and Teaching Collections (symbolic code: TCWC), or Museum of Southwestern Biology (MSB). Additionally, toe and/or tail tips were collected non-destructively from animals caught in the field that were subsequently released. All tissue samples were stored in 95% EtOH. Whole genomic DNA was extracted from tissues using the DNeasy Blood and Tissue kit (Qiagen).

**Figure 1.**
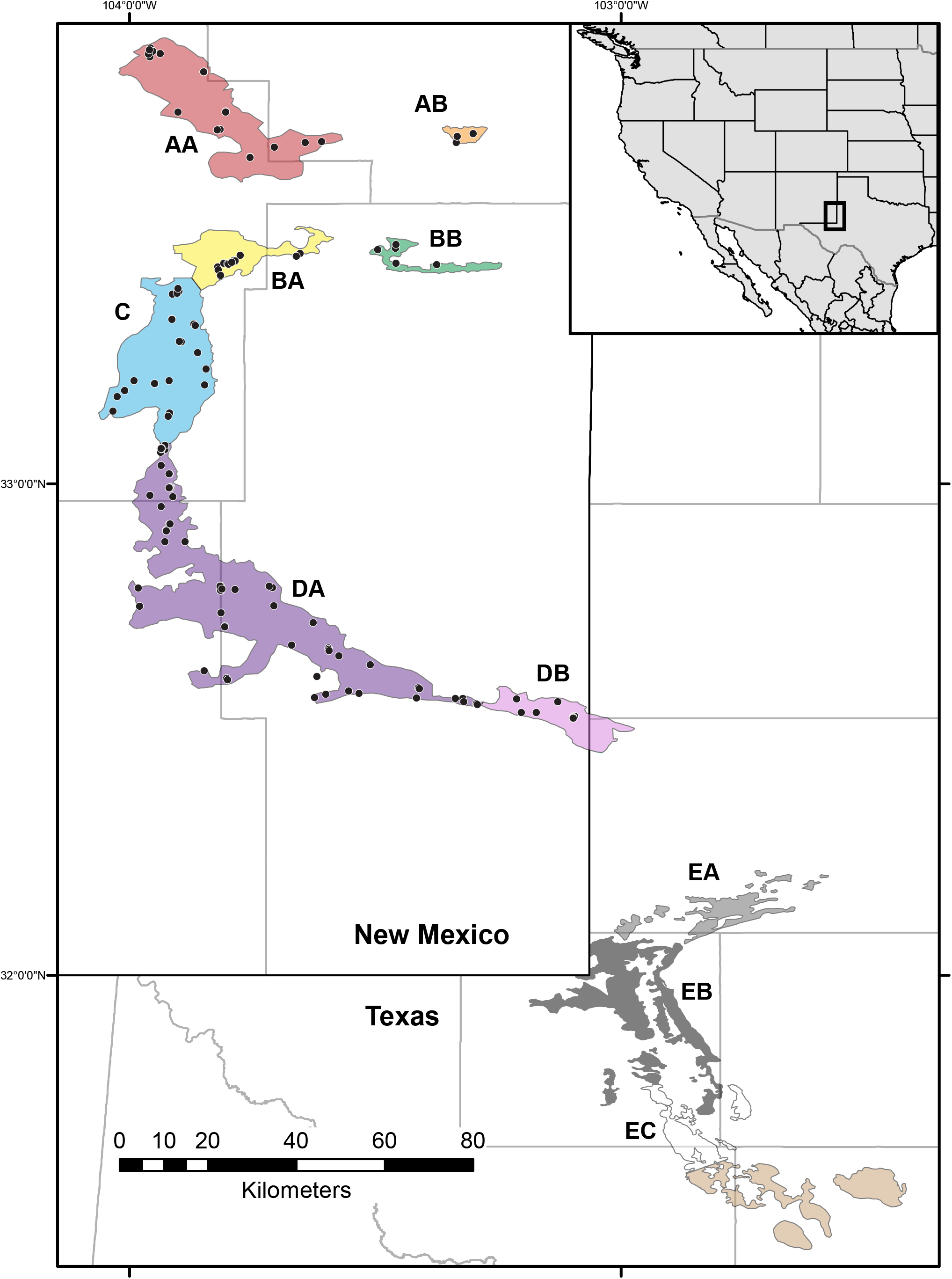
Collection localities for samples of *S. arenicolus* from New Mexico included in this study and the outline of the species’ range and suitable shinnery oak-sand dune habitat (from Laurencio and Fitzgerald, 2010). Specific localities are not shown for Texas due to legal confidentiality agreements with landowners. Colored portions of the species’ range correspond to phylogroups and geographic regions referred to in text. Brown indicates potential habitat in Texas where *S. arenicolus* has not been found; one locality exists in this region from 1970. Presence/absence data and habitat suitability maps could be used to more precisely delineate geographic boundaries of the phylogroups within areas of suitable habitat.

### DNA sequence data

We targeted two mitochondrial and four nuclear loci for DNA sequencing. PCR amplification of the mitochondrial loci NADH-dehydrogenase 1 (*ND1*) and cytochrome-b (*cyt-b*) and two protein coding nuclear loci prolactin receptor (*PRLR*) and *R35* used previously published primers (Irwin et al., 1991; Leaché and McGuire, 2006; Leaché, 2010). We used two additional anonymous nuclear loci designed from a genomic library enriched for microsatellite repeats: *sarANL298* (scar298anl.F: 5’-ATGGGAAGGCTTAAAATGAATC; scar298anl.R: 5’-TGTGACTTAGGGAACTGGGTATGT) and *sarANL*875 (scar875anl.F 5’-CTTACCATTCAACCCTTCCTTG; scar875anl.R 5’-CTAGAGCAGACCAGTTCAATGTAAT). All PCR were conducted in 10 μl total volume. Annealing temperature for the new nuclear loci was 54°C.

We used 0.4 μl ExoSAP-IT (USB/Affymetrix) and 1.6 μl water to clean 5 μl of PCR product. One μl of clean PCR template was used in each cyclo-sequencing reaction using the same locus-specific primers used in amplification. Sequencing reactions were cleaned and run on an ABI 3730xl at the Duke Sequencing Facility or the Biotechnology Resources Center of Cornell University. Chromatograms were verified and cleaned in Geneious R9 (https://www.geneious.com). Heterozygous sites in nuclear sequences were called with the appropriate ambiguity code. Sequences at each locus were aligned using the MAFFT (Katoh, 2005) plug-in in Geneious. All sequence data will be submitted to GenBank.

Because the mitochondrion is inherited as a single unit without recombination, we concatenated the two loci (ND1 and Cyt-*b*) into a single alignment. Each of the four nuclear loci were treated independently. All sequences at each locus were aligned in Geneious and alleles at nuclear loci were determined using the program PHASE (Stephens et al., 2001) and the helper program SeqPHASE (Flot, 2010).

### Microsatellite genotype data

Nuclear microsatellite loci were developed from a 454-library enriched for microsatellite motifs developed at Cornell University Evolutionary Genetics Core Facility. After initial screening of loci, we used the Qiagen Type-It microsatellite PCR kit to genotype individuals at these loci in five multiplex reactions (Supp. Mat. Table 1). Forward primers for all loci were tagged with a fluorescent dye and samples were genotyped on an ABI3730xl at the Biotechnology Resource Center of Cornell University with GeneScan 500 LIZ size standard (Thermo Scientific). Alleles were called and verified for all individuals using GeneMarker 2.6. Prior to subsequent genetic analyses, all variable loci were tested for the presence of null alleles and selection by testing for Hardy-Weinberg Equilibrium (HWE) and for evidence of linkage disequilibrium using GenePop (Rousset, 2007). The final dataset included genotypes for all individuals at 27 variable and neutrally evolving nuclear microsatellite loci.

### Data analysis

#### Summary statistics

We used PAUP (Swofford, 2002) to determine the number of parsimony informative sites for each sequence alignment and DNAsp v6 (Rozas et al., 2017) to calculate the number of unique haplotypes, the number of segregating sites (S), nucleotide diversity (π), and the average number of nucleotide differences (k) for each sequence alignment.

#### Haplotype networks

We constructed parsimony networks in TCS (Clement et al., 2000) for complete mtDNA haplotypes for *S. arenicolus*. Because the results generated by network methods can be strongly influenced by missing data (Joly et al., 2007), we first omitted all individuals with missing sequence data for one of the two mitochondrial loci. We additionally omitted individuals for which we did not have locality information. The final haplotype network for mtDNA contained 195 individuals. We additionally constructed parsimony networks for the phased alleles at each nuclear locus.

#### Phylogenetic analysis

For the mitochondrial DNA, we estimated the phylogenetic relationships among *S. arenicolus* under both maximum likelihood and Bayesian frameworks. *Urosaurus ornatus, Uta stansburiana, Phrynosoma coronatum, Sceloporus jarrovii, S. merriami, S. occidentalis*, and nine individuals of *S. graciosus* were used as outgroups (following Chan et al., 2013). Concatenated mtDNA alignments were first reduced to unique sequences using a Python script from BioPython (sequence_cleaner.py). We estimated the best-fit model of sequence evolution at each codon position of each gene in DT-ModSel (Minin et al., 2003) and partitioned phylogenetic analyses by gene and codon position. The best fit models by DT-ModSel were a SYM+G for the first codon position of each gene, HKY+I for the second codon position of each gene, and TrN + I + G and TrN + G for the third codon position of *ND1* and *Cyt-b* respectively. We estimated the phylogeny under a Bayesian framework in MrBayes v3.2.6 (Ronquist and Huelsenbeck, 2003; Ronquist et al., 2012) excluding individuals with missing data. TrN models were expanded to GTR for Bayesian analyses and the final analysis consisted of two independent runs each of 50 million generations sampled every 5,000 generations. All parameters were checked for adequate mixing and convergence, and the maximum clade credibility tree was summarized in MrBayes.

#### Population genetic analysis

We estimated within population diversity and among population pairwise F_ST_ for mtDNA as well as microsatellite data assuming membership to the phylogroups based on the Bayesian phylogeny. Estimates of F_ST_ were done in Arlequin (Excoffier and Lischer, 2010) for mtDNA and in FSTAT for microsatellite data. Because samples were distributed evenly throughout the range of *S. arenicolus*, we additionally conducted population genetic analyses without any assumption of population membership using assignment methods in Structure 2.3.4 (Pritchard et al., 2000). In Structure, we tested assignment of all individuals to *K* populations from K=1 to 10. At each K we conducted 10 replicate runs each consisting of 1 million generations with the first 50% discarded as burn-in. We used StructureHarvester (Earl and vonHoldt, 2011) to examine all runs and CLUMPP (Jakobsson and Rosenberg, 2007) and DISTRUCT (Rosenberg, 2004) to visualize population membership. Structure runs with all individuals supported K=2, so subsequent runs investigated further partitioning with each major group. For each subset of data, we tested K=1 to 5 each with 10 replicate runs at each K each consisting of 2 million runs with the first 50% discarded as burn-in.

#### Demographic analyses

We estimated the historical demographics for each of five primary phylogeographic regions (A-E) identified in mtDNA analyses. We used multilocus sequence data to construct extended Bayesian skyline plots in BEAST 2.5.0 (Drummond et al., 2005, 2012; Heled and Drummond, 2008). Each dataset included concatenated mtDNA alignments in addition to phased genotypes for each of the four nuclear loci. Substitution models for each locus were set based on MrModeltest (Nylander, 2004; Supp. Mat. Table 2). All runs assumed a relaxed molecular clock with a log-normal distribution for the mtDNA partition and strict molecular clocks for the nuclear partitions. The rate for mtDNA was set with a log normal distribution with mean of 1×10^−8^ substitutions/site/year and SD of 0.27 following (Chan et al., 2013). Parameter trends were examined in Tracer to check for adequate mixing within runs and convergence across runs. Final runs were 50 million steps sampled every 5,000 steps for regions B, C, and E. The final runs for regions A and D were 100 and 200 million steps sampled every 10,000 and 20,000 steps, respectively. Extended Bayesian skyline plots we generated after discarding the first 25% sampled steps as burn-in.

#### Hypothesis testing

Based on the results of phylogenetic analyses and assignment tests, we tested three alternative hypotheses of divergence and population expansion among three geographic groups (Northern Mescalero Sands, Southern Mescalero Sands, and Monahans Sandhills) assuming that the Monahans Sandhills populations were ancestral and of constant population size (Chan et al., 2009; Supp. Mat. Figure 1). We used approximate Bayesian computation to evaluate support for these models and estimate demographic parameters of the best supported model in DIYABC (Cornuet et al., 2008). Analyses included mtDNA and phased nuclear sequences. Locus parameters were specified after estimation of substitution models for each locus in DT-ModSel. The prior for the mtDNA mutation rate was set as a normal distribution with a mean of 1×10^−8^ substitutions per site per year and nuclear substitution rates were set as uniform distributions. Initial runs were used to determine adequate priors for demographic parameters. The final analysis included 2 million samples for each divergence model (6 million total) with a linear regression step to extract the closest 1% of samples and determine the best supported model of the three. For the best supported model, we used the same selection/rejection process to estimate divergence times and demographic parameters from the closest 1% of the 2 million samples.

## Results

### Summary statistics

Sample sizes, alignment lengths, the number of unique haplotypes, number of segregating sites, average nucleotide differences, and nucleotide diversity are reported in Supp. Mat. Table 3. As expected, nuclear loci were less variable than mtDNA though nucleotide diversity was similar for mtDNA and two nuclear loci. We also recovered multilocus genotypes for 237 individuals at 27 microsatellite loci that conformed to HWE expectations and did not show any evidence of linkage or null alleles. The average number of alleles per locus was 16.4 with a range from 3 to 35 (Supp. Mat. Table 4).

### Haplotype networks and phylogenetics

Mitochondrial haplotype networks revealed geographically associated haplotype groups for mtDNA that largely correspond to regions of grossly contiguous habitat (Figure 2). In the Northern Mescalero Sands, there are three main haplotype groups corresponding largely with the A regions (Figure 1; AA and AB), the B regions (BA and BB), and the C region, though the genetic divergence among these three groups is small. Common haplotypes are shared across regions, but derived haplotypes are unique to each region. Regions AB and BB have genetic diversity that is primarily a subset of the diversity found in AA and BA, respectively. The Southern Mescalero Sands (Regions DA and DB) are genetically divergent from the Northern Mescalero Sands populations with the barrier between the two groups reflecting a west-east constriction in the distribution of potentially suitable habitat (referred hereafter as “the Skinny Zone”). Among the Southern Mescalero Sands individuals in region DA, we find a single widespread haplotype and multiple derived haplotypes. In addition, region DB at southernmost tip of the Southern Mescalero Sands contains a cluster of derived haplotypes.

**Figure 2.**
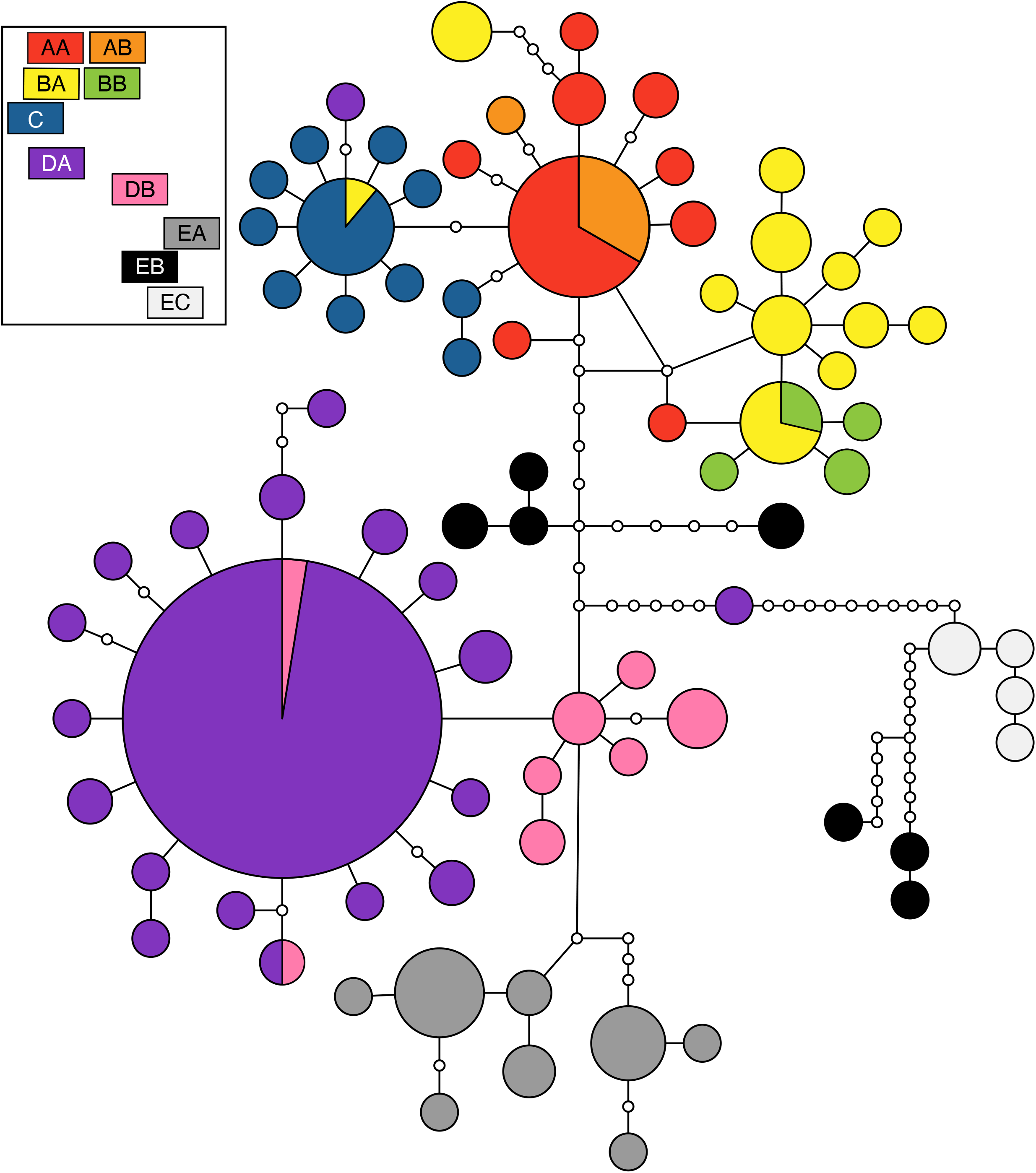
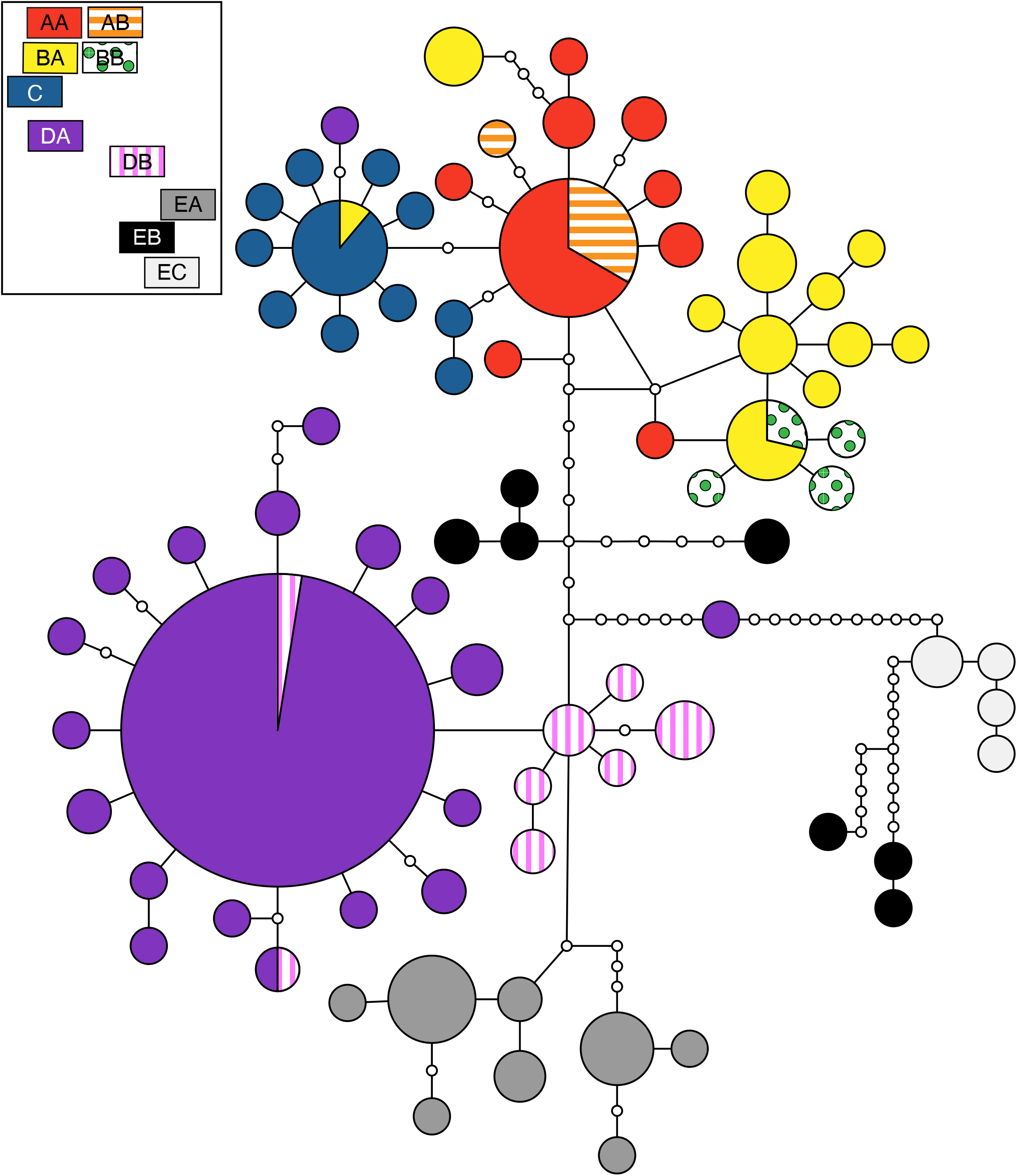
Haplotype networks based on concatenated mtDNA sequences. Circles represent unique haplotypes with the size of the circle corresponding to the relative abundance and the color referring to the region of origin of individuals with that haplotype (see boxes in upper left representing geographic approximations of each region). Lines connecting haplotypes represent one mutational step. Small white circles represent unsampled haplotypes. [Alternate version for individuals with color vision deficiencies included].

Populations in the Monahans Sandhills are genetically distinct from all other *S. arenicolus* populations, but do not form a single haplotype group. There is high sequence divergence among haplotypes from the Monahans Sandhills despite occurring in a relatively restricted geographic area and they are distantly related to Mescalero Sands haplotypes. The EA and EC areas each have unique haplotypes without a single, most common haplotype. The EB haplotypes fall out into two main groups, one that is equally distant to northern and southern Mescalero Sands haplotypes and one that is distantly related to all other recovered haplotypes (Figure 2).

In general, nuclear gene regions had much lower genetic diversity with very little genetic structure (Supp. Mat. Figure 2). Across all four nuclear loci, we found a similar pattern with the most common haplotypes occurring in most, or all regions. At PRLR and scar875, several derived loci were unique to Monahans Sandhills populations and Monahans Sandhills plus Southern Mescalero Sands populations. With one exception (AB locus R35) regions AA, AB, and BB did not have any unique nuclear alleles.

Phylogenetic reconstructions largely corroborated the groups found in the network analyses (Figure 3). We recover *S. arenicolus* as monophyletic (PP = 1). Monahans Sandhills populations were paraphyletic with respect to Mescalero Sands populations with the southern-most Monahans Sandhills individuals forming a weakly supported clade (PP = 0.8657) sister to all other *S. arenicolus*. Among the remaining individuals, there is strong support for a Northern Mescalero Sands clade including individuals north of the skinny zone (PP = 0.9758) and moderate support for a Southern Mescalero Sands – Monahans Sandhills clade that includes individuals south of the skinny zone and the northern and central Monahans Sandhills (PP = 0.9345). Within the Northern Mescalero Sands clade, we recover support for some clusters of individuals, but do not find well-supported clades corresponding to distinct geographic regions. Individuals from region A, at the northern end of the range, form a basal polytomy relative to otherwise well-supported clades containing most individuals from regions B and C, and several from region A and one from D (TCWC 94831). Support for a clade that includes most B individuals is high (PP = 0.9639) as is support for two different clades that each primarily include individuals from C (PP = 1). We do not recover very much genetic resolution for individuals south of the skinny zone in the Southern Mescalero Sands or the northern or central Monahans Sandhills. Notably, individuals from Monahans Sandhills are paraphyletic and their relationships largely unresolved. While most individuals from regions cluster with other individuals from the same region, as expected from the haplotype network, there are a few individuals that fall out with individuals from different regions.

**Figure 3.**
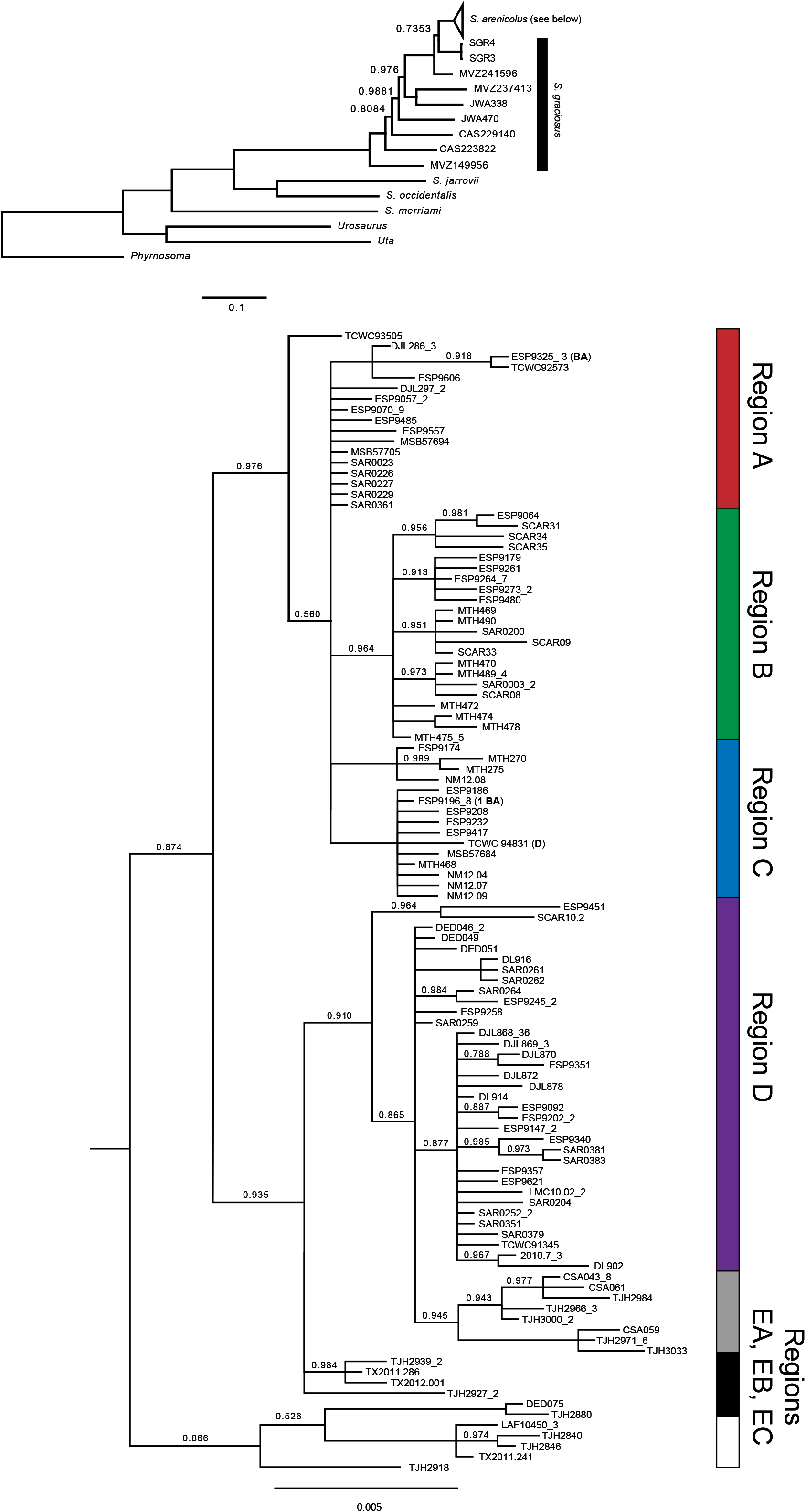
Majority-rules consensus tree from Bayesian phylogenetic analysis of concatenated mtDNA sequence data. Posterior probability for all nodes is 1 unless otherwise indicated. Tips are labeled with a sample name followed by the number of samples with an identical haplotype. Regions of collection are indicated vertically with several exceptions listed parenthetically in terminal name.

### Demographic estimates

Extended Bayesian skyline plots match the inferences made from the haplotype networks (Supp. Mat. Figure 3). We see evidence of recent population expansion in region D and demographic stability in region E. Regions A, B, and C show some evidence of population expansion though the credible intervals around the most recent population sizes is large and does not exclude the possibility of demographic stability.

### Population Genetics

Pairwise F_ST_ among populations was high among regions with values significantly different from zero ranging from 0.099 to 0.904 for mtDNA and 0.026 to 0.236 based on microsatellite loci (Table 1).

**Table 1.** Pairwise F_ST_ values among regions for microsatellite genotypes (above diagonal) and mitochondrial sequence data (below diagonal). Values significantly different from zero (at alpha < 0.05) are indicated in bold. The significance of some values was not able to be determined because of low genetic variability, indicated with italics.

|  | AA | AB | BA | BB | C | DA | DB | EA | EB | EC |
| --- | --- | --- | --- | --- | --- | --- | --- | --- | --- | --- |
| AA |  | **0.0807** | **0.0349** | *0.0689* | **0.0358** | **0.1443** | **0.1723** | **0.1488** | **0.1654** | **0.1457** |
| AB | 0.0279 |  | 0.1077 | *0.1911* | 0.0749 | **0.1988** | 0.2374 | 0.2080 | 0.2356 | 0.2010 |
| BA | **0.3389** | **0.3151** |  | *0.0512* | **0.0264** | **0.1337** | **0.1594** | **0.1349** | **0.1388** | **0.1033** |
| BB | **0.6593** | **0.7667** | **0.2147** |  | *0.0807* | *0.1643* | *0.2116* | *0.1688* | *0.1830* | *0.1592* |
| C | **0.4102** | **0.3688** | **0.4605** | **0.6583** |  | **0.1271** | **0.1523** | **0.1213** | **0.1350** | **0.1071** |
| DA | **0.8466** | **0.8584** | **0.8083** | **0.8687** | **0.8503** |  | **0.0411** | **0.0882** | **0.1188** | **0.0990** |
| DB | **0.8381** | **0.8395** | **0.7483** | **0.8439** | **0.8173** | **0.3311** |  | **0.1213** | **0.1554** | 0.1302 |
| EA | **0.7984** | **0.7723** | **0.7372** | **0.7842** | **0.7858** | **0.5767** | **0.4206** |  | **0.1188** | **0.1017** |
| EB | **0.5554** | **0.4000** | **0.5021** | **0.4513** | **0.5523** | **0.6668** | **0.4663** | **0.5302** |  | 0.0701 |
| EC | **0.8406** | **0.8125** | **0.7540** | **0.8216** | **0.8099** | **0.8872** | **0.8424** | **0.8108** | **0.4119** |  |

Assignment tests based on microsatellite data reveal nested structure at multiple spatial scales (Figure 4). Across all samples, our analyses recover two groups with some admixture. The geographic break between these two groups corresponded to the Skinny Zone of the Mescalero Sands with some individuals in this area being admixed. Further assignment tests in Structure with nested subsets of the data indicate that these admixed individuals are aligned with individuals in the Southern Mescalero Sands. We recover distinctive groups in the Northern Mescalero Sands with some admixture as well. Region A individuals are distinct from region B + C individuals although, assignment plots suggest some admixture between western A populations (AA) and populations in region C, corroborating results from the mtDNA haplotype networks. Analysis of the AA-AB groups recover AB as genetically distinct corroborating F_ST_ estimates (Table 1). Analysis of the BA-BB-C group supports BB and C as distinct from one another with BA having genetic affinities to both. Together these results show a clear genetic break between the Southern Mescalero Sands and Monahans Sandhills populations. For the Southern Mescalero Sands populations there is an additional genetic break between regions DA and DB coinciding with another constriction in suitable habitat. Among Monahans Sandhills samples, EA individuals are largely distinct from EB+EC. Individuals from EB and EC are somewhat distinct from one another though not all individuals within a region cluster unambiguously with others in the group.

**Figure 4.**
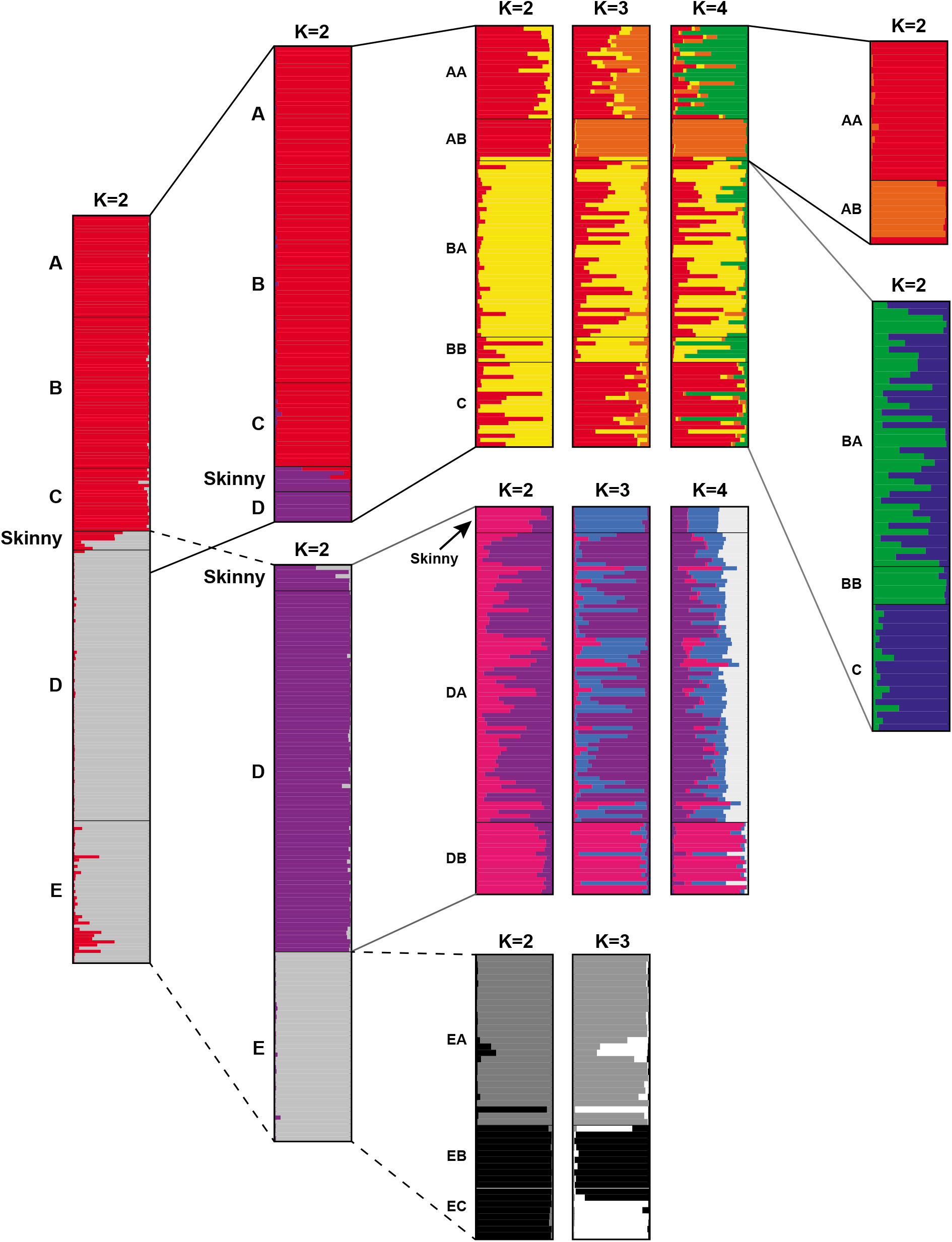
Individual assignment plots from nested Bayesian assignment tests in STRUCTURE. Results at alternate values of K are shown for some subsets of individuals.

### Hypothesis testing

We recover strongest support (PP = 0.9991) for a divergence scenario that involved colonization of the Northern Mescalero Sands from Monahans Sandhills populations around 34.8 Kya (CI 17.7-108 Kya) followed by colonization of the Southern Mescalero Sands from Monahans Sandhills populations more recently, around 16.3 Kya (CI 7.9-41 Kya; Figure 5). It is possible that the initial colonization of Northern Mescalero Sands included colonization of the Southern Mescalero Sands, with subsequent local extinction and recolonization, or genetic replacement. It is important to note that the 95% credible intervals for all estimates of divergence time and population size are broad. In fact, though the time of expansion in the Northern Mescalero Sands (T_exp1_)was constrained in individual ABC simulations to occur after the divergence between the Northern Mescalero Sands and the other regions (T_anc_), the median estimate for the former T_exp1_ precedes the median divergence time, T_anc_ (Figure 5), though both estimates have extremely broad and overlapping credible intervals.

**Figure 5.**
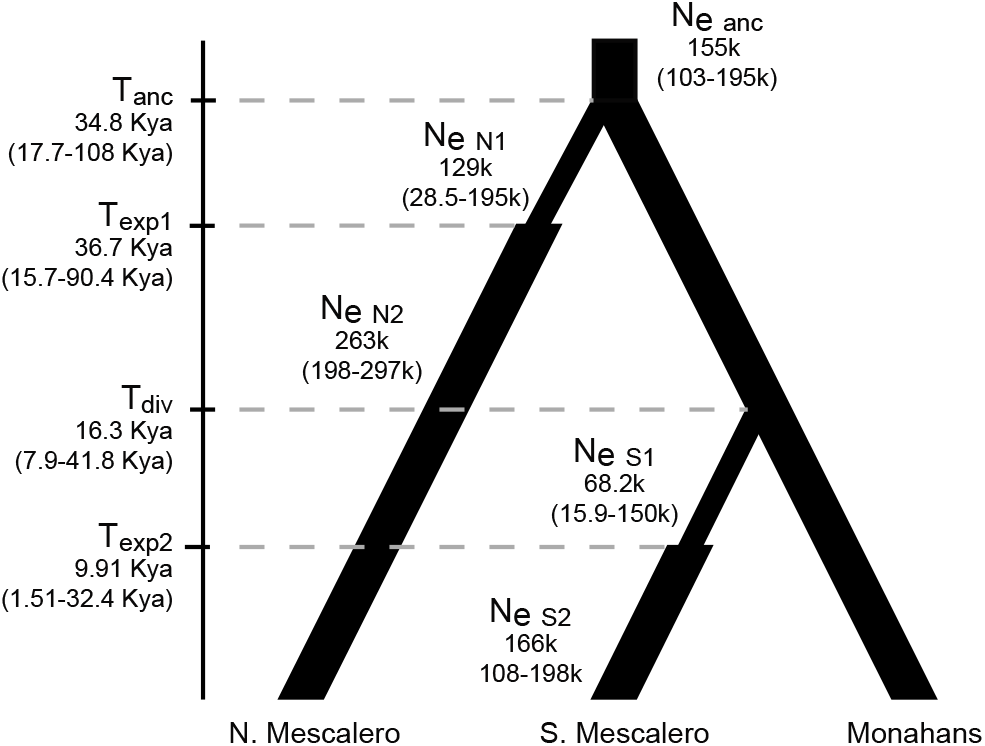
Estimates of divergence and expansion times as well as current and historical effective population sizes for the best supported model from ABC analysis of the complete genetic dataset.

## Discussion

Sampling of *S. arenicolus* throughout the entire range provides greater resolution of the evolutionary patterns of divergence of this narrowly distributed habitat specialist. We find support for multiple genetic groups within *S. arenicolus* suggesting limited migration in this habitat specialist. In particular, we find genetic structure beyond the three mitochondrial groups described in Chan et al. (2009). Patterns of divergence recovered by mtDNA corroborate nuclear microsatellite data and demonstrate the importance of the landscape-scale configuration of areas of habitat on the phylogeographic structure of this habitat specialist. With thorough geographic sampling we are able to identify regions that have served as barriers to population connectivity and characterize historical demographics across evolutionary time scales.

Lineages of *S. arenicolus* in the Mescalero Sands and Monahans Sandhills have independent and distinct histories that are associated with the timing of sand deposition and dune formation in these sub-regions. Indeed, the Mescalero Sands and Monahans Sandhills have related, but distinguishable geologic histories (Muhs and Holliday, 2001; Rich and Stokes, 2011; Muhs, 2017). Both Mescalero Sands and Monahans Sandhills are sand sheets of the Southern High Plains deposited over older, compact eolian deposits comprising the Black Water Draw formation (204 - 43 Kya; Rich and Stokes, 2011). Sand accumulation and dune formation has occurred repeatedly with current sand sheet age estimates for Mescalero and Monahans as 29.2 and 22.2 Kya respectively and with a more recent deposition ~ 7.5 Kya (Rich and Stokes, 2011). Though *S. arenicolus* sampled from the Monahans Sandhills do not form a monophyletic group, it is clear that they are distinct from Mescalero Sands populations with estimated initial divergence between these two regions occurring long ago (34.8 Kya, CI 108-17.7 Kya; Figure 5). While the estimated divergence is older than the estimated age of the most recent sand deposition, this is a dynamic landscape that has undergone cycles of sand deposition during periods of aridity (Holliday, 1989; Rich and Stokes, 2011) such that this divergence is most likely associated with previous episodes of sedimentation and dune formation. There are broad CI around estimates of divergence and population expansion, but these estimates generally coincide with the sand age of Northern Mescalero Sands. Furthermore, the estimate of the most recent deposition falls within the CI for colonization and expansion times for the Southern Mescalero Sands.

The location and movement of sand dune formations has changed over millennia (Muhs and Holliday, 1995, 2001; Muhs, 2017). While the presence of sand dunes alone does not indicate the presence of shinnery oak-sand dune ecosystem, the distribution of habitat suitable to *S. arenicolus* has likely shifted in its occurrence and connectivity over geologic time. Given the dynamic nature of the landscape to which *S. arenicolus* is endemic, it stands to reason that dynamic histories also characterize the phylogeographic and population genetic structure in this species. Though the age of the Mescalero Sands and Monahans Sandhills geologic formations are uncertain, our data suggest that Monahans Sandhills was the source population from which Mescalero Sands *S. arenicolus* populations were colonized. Mescalero Sands populations, which lie to the north of the Monahans Sandhills, are comprised of at least two distinct lineages, but nuclear microsatellite data and ABC analyses suggest that the Southern Mescalero Sands populations are more closely related to the Monahans Sandhills populations than to northern Mescalero Sands populations. The two sand formations are not currently connected by suitable habitat (Figure 1), but presumably were connected in the past facilitating the colonization of Mescalero Sands from Monahans Sandhills. by *S. arenicolus*. The current range map (Figure 1) is informed by currently occupied habitat, but given the dynamic history of the shinnery oak-sand dune ecosystem, potentially suitable habitat connecting regions may have occurred in the past. The configuration of available habitat is varies across time which presumably causes concordant shifts in species’ distributions.

Our genetic data suggest that the colonization event associated with the current Southern Mescalero Sands populations occurred separately from the event that resulted in the Northern Mescalero Sands populations. Colonization of the Northern Mescalero Sands and divergence from the Monahans Sandhill source population is estimated to have occurred approximately 34 Kya followed by population expansion (Figure 5; Supp. Mat. Figure 3). While recognizing that there are broad confidence intervals around the estimated time of this event, it is plausible that this divergence was associated with the deposition of loose aeolian sands over the Blackwater Draw Formation (Rich and Stokes, 2011). The second divergence was between the southern Mescalero Sands and Monahans Sandhills occurring later, around 16.3 Kya. This is similar to the age of more recent sand deposits in the Mescalero Sands, and subsequent population expansion with a median estimate of 9.9 Kya coincides roughly with the ages of the most recent aeolian deposits. This result suggests that after the colonization of the Mescalero Sands 34 Kya by *S. arenicolus*, habitat between the Mescalero Sands and Monahans Sandhills contracted or that Southern Mescalero Sands populations became extirpated and this area was later recolonized. Both scenarios seem plausible given what we know about *S. arenicolus* ecology and the dynamic nature of this system.

*Sceloporus arenicolus* requires interconnected shinnery-oak blowouts to support populations (Ryberg et al., 2013; Leavitt and Fitzgerald, 2013). Shinnery oak flats or isolated dune blowouts impede movements and isolate populations. The divergences that we see across the Mescalero Sands and Monahans Sandhills correspond largely with the geographic extent of potentially suitable habitat identified in several studies of *S. arenicolus* (Fitzgerald et al., 1997; Laurencio and Fitzgerald, 2010; Walkup et al., 2018). We are also able to reconstruct historical population demography and recover variable, and sometimes dynamic, histories across populations of *S. arenicolus*. For instance, we find support for a major genetic break that coincides with the Skinny Zone, a narrow constriction (~ 3 km wide) in the central Mescalero Sands. This narrow zone of habitat for *S. arenicolus* is indicative of a long-standing barrier to dispersal and is now a point of secondary contact between divergent Northern and Southern Mescalero Sands populations.

We find shallow divergence, but distinct genetic diversity among the Northern Mescalero Sands regions indicating that habitat suitability also impacts population genetic connectivity at these finer spatial scales. Populations in some of these regions, like AB and BB, have diverged in isolation, suggesting a founder effect in line with the major direction of sand dune movement (Muhs, 2017). We additionally confirm a recent colonization and subsequent rapid population expansion in the southern portion of the Mescalero Sands (Region D). Finally, among the Monahans Sandhills samples we documented highly divergent alleles, deep divergence among populations, and relative population stability. We recovered at least three divergent groups among the Monahans Sandhills individuals indicating limited movement among older populations retaining ancestral genetic diversity.

The historical demography and patterns of divergence are reflected in the microsatellite data as well as the more slowly evolving mtDNA sequence data indicating that population structure is the result of longstanding habitat dynamics and restrictions to gene flow at multiple spatial scales, not just more recent anthropogenic change. Demographic studies of *S. arenicolus* have emphasized the importance of a network of suitable habitat at multiple spatial scales to support metapopulation dynamics and population persistence (Ryberg et al., 2014). Landscape-ecological analyses of presence and absence of lizard community membership across the Mescalero Sands demonstrated that landscape heterogeneity, not dispersal, explained community assembly and meta-community structure (Ryberg and Fitzgerald, 2015, 2016). The occurrence of the habitat specialist *S. arenicolus* was a driver of this pattern. As such, because the fine-scale distribution of suitable habitat is critical for local presence of *S. arenicolus*, we suggest the composition and configuration of the landscape with respect to unsuitable habitat types determines patterns of genetic connectivity across the range. The divergences we detect reinforce that extensive habitat may be necessary to support gene flow among populations and that habitat quality and habitat configuration at finer scales may be of critical importance to identifying potential corridors. Importantly, it is clear that the shinnery oak-sand dune ecosystem is a dynamic landscape where the configuration of habitat patches can change over decades and millennia. We know from phylogeographic studies that specialists in changing environments undergo repeated episodes of isolation and divergence (Roderick et al., 2012). While it is impossible to reconstruct the specific, chronological habitat configuration for the Mescalero Sands and Monahans Sandhills, it is likely that networks of suitable habitat have diverged and coalesced repeatedly over time (e.g., Dzialak et al., 2013). Source-sink dynamics are important at local and contemporary spatial and temporal scales (Ryberg et al., 2013; Walkup et al., 2019), and this may translate to evolutionary patterns of population genetic structure at broader spatial scales and longer time scales. Under this model, habitat patches shift in their extent and distribution over time due to geological processes. The divergence and coalescence of habitat patches across time results in repeated local extinction, population divergence, and recolonization. In support of this scenario, we find population genetic and demographic patterns that reflect such dynamic processes and their variability across the landscape. For instance, the Southern Mescalero Sands is a more rapidly shifting sand dune formation (Muhs and Holliday, 1995, 2001; Muhs, 2017) in comparison to the sand sheets of the Monahans Sandhills formation which are more stable and less dynamic (Machenberg, 1984). The Southern Mescalero Sands may be characterized by local extinction and recolonization whereas the slower movement of the Monahans Sandhills may maintain demographically stable and isolated populations over longer time periods.

The patterns of divergence and gene flow that we see in *S. arenicolus* are not surprising of a habitat specialist inhabiting a dynamic landscape. Based on demographic studies (Leavitt and Fitzgerald, 2013; Walkup et al., 2019) and observations of *S. arenicolus* (Ryberg et al., 2012; Leavitt and Acre, 2014; Walkup et al., 2018, p. 2018; Young et al., 2018), individuals do not move large distances. Their strict habitat requirements, and the naturally patchy and temporally dynamic qualities of this habitat, suggests that populations should be subdivided. The nestedness of genetic structure in *S. arenicolus* mirrors the hierarchical nature of their habitat preference: individuals require suitable blowouts within a matrix of shinnery oak, and populations are supported by a network of connected shinnery oak-sand dune complexes. While the genetic consequences of metapopulation dynamics have typically been explored at fine spatial and temporal scales, our phylogeographic study shows that these metapopulation dynamics may also leave their signature at broader spatial scales, in this case, across the range of this endemic lizard.

## Conservation

The shinnery oak-sand dune habitats of Mescalero Sands and Monahans Sandhills have experienced severe habitat degradation and fragmentation, particularly in the southern portions of the range of *S. arenicolus* (Leavitt and Fitzgerald, 2013; Walkup et al., 2017). Recent ongoing fragmentation due to human activities (e.g. highways and caliche roads built for oil field development) is known to decrease connectivity among populations and interrupt metapopulation dynamics leading to extinction of local populations (Ryberg et al., 2013, 2014; Leavitt and Fitzgerald, 2013; Walkup et al., 2017). Fragmentation of the shinnery oak-sand dune ecosystem increases the likelihood that ancestral diversity and unique evolutionary lineages will be lost. Our findings highlight regions to be considered as genetically distinctive conservation units as well as underscore the unique genetic and demographic history of different regions within the range of *S. arenicolus*.

## Acknowledgments

Thank you to J. T. Giermakowski (MSB), R. Macey, C. Spencer (MVZ), and J. Vindum (CAS) for access to tissue samples. We thank the many research assistants who helped with tissue collection, DNA sequencing, and genotyping, including J. Moberg, E. Gibson, D. Cavero, and T. Caspi. Sequencing was conducted at the Genome Sequencing & Analysis Core Resource of the Duke Institute for Genome Sciences and Policy. Genotyping of microsatellite loci was completed at the Biotechnology Core Facility of Cornell University. Portions of this research were provided by the Dunes Sagebrush Lizard/ Lesser Prairie Chicken Candidate Conservation Agreement (CCA) Research Fund administered by CEHMM, Carlsbad, New Mexico, and by USA Bureau of Land Management, and by the State of Texas Comptroller of Public Accounts. This is contribution number xxxx of the Biodiversity Research and Teaching Collections, Texas A&M University.

