## Supplementary Materials for "Phylogeographic structure of the dunes sagebrush lizard, an endemic habitat specialist"

Supp. Mat., Table 1. Microsatellite locus information.

| Multiplex | Locus Name | Dye | Repeat | Size | Primer Sequences (5’-3’) | Notes |
|---|---|---|---|---|---|---|
| Multiplex 1 | sarms0344 | VIC | $(ACT)^{11}$ | 250 | F: CTGAGGACTTTGGTTTGAGGAA<br>R: ACCCAGTATGAGAGGAATGAAGC | |
| 57°C annealing | sarms0473 | PET | $(TG)^{13}$-T-$(TG)^{3}$ | 155 | F: CCTCATCTGTATCCCTCTCATTAG<br>R: AGCCAGTCATCTTCTCTTTCATAC | |
| Q solution | sarms3490 | NED | $(ATC)^{6}$ | 182 | F: TTGCTTGTAACACCCACTGATAG<br>R: CCTGTACACCGCTATGATCAATG | |
| | sarms5185 | 6-FAM | $(GAT)^{5}$ GGT $(GAT)^{7}$ | 230 | F: GCCAATGCAAGACAGAAAATAGAA<br>R: GGGAGGAGGGTTGGGATGC | |
| | sarms5839 | NED | $(CTAT)^{15}$ | 335 | F: ATGGCAGCTTTCTTTCTTG<br>R: TGTTATAGTCAGTGGGTTGGTA | |
| | sarms6346 | PET | $(AGAT)^{11}$ | 369 | F: ATGACTAATATCCACCCACCCTG<br>R: TTCAGCATGCTTGTAAGAACCTG | |
| Multiplex 2 | sarms0506 | 6-FAM | $(TTG)^{6}$ | 206 | F: TCAGGAAAAGGCGGGGTAT<br>R: GCAGGGAAAAACAGAGCAGAA | |
| 60°C annealing | sarms2620 | NED | $(AAGG)^{12}$ | 354 | F: ACAACAGCATTTAGGAAGGAAGG<br>R: AAACTCCTACCTTCCTCGAACG | Failed HWE |
| Q solution | sarms2770 | PET | $(AAC)^{8}(AC)^{3}$ | 190 | F: TGGGTCACATACTCTCAGC<br>R: GCCATGCAGGAATACACT | |
| | sarms3645 | 6-FAM | $(GATA)^{18}$ | 408 | F: ACTAGGTCTTCTTGTTCATCATTG<br>R: GGGCGGTACTCATCTCTGTTCCTA | |
| | sarms4015 | PET | $(CTTT)^{12}$ | 421 | F: CCCAGCCCAAAAGGAACC<br>R: CTCAGAGGCATCAAGAAGACA | |
| | sarms4354 | NED | $(CA)^{18}$ | 160 | F: TCCCTGATCACCCCTCTGC<br>R: TTGCTGAACGGTGATGTCTTG | |
| | sarms6064 | VIC | $(AGT)^{7}$ ACT $(AGT)^{5}$ | 251 | F: CAAGAACAGGGGGATGACTA<br>R: CAAGGGCAATGAACTCTGGA | |
| Multiplex 3 | sarms0739 | PET | $(GT)^{10}$ | 163 | F: GCAAAATGGAGAATCGTGTGAGTA<br>R: TAGTGGGGAATAGGAAGGGTAGG | |
| 58°C annealing | sarms4884 | 6-FAM | $(TG)^{15}$ | 160 | F: GTTTCCTTCATCTGTATCCTCTCA<br>R: AGCCAGTCATCTTCTCTTTCATAC | Failed HWE |
| no Q solution | sarms5968 | 6-FAM | $(ACT)^{11}$ | 270 | F: CAAGGGCAATGAACTCTGGA<br>R: GTGGTGGCGGCTCTGTG | |
| | sarms2196 | VIC | $(AC)^{6}$ | 152 | F: GATGATAGCTCCTTCAGTTGGTG<br>R: AGCAAATCATAGCACCACAGAAG | Monomorphic |
| | sarms7111 | PET | $(TCTT)^{13}$ | 390 | F: AACAAATGCCCCACC<br>R: GGCAAGACCCAAACACTA | |

| | | | | | | |
|---|---|---|---|---|---|---|
| Multiplex 4 | sa60.12 | PET | $(GT)^{24}$ | 122-148 | From Chan et al. 2007 | |
| 58°C annealing | sarms0001 | PET | $(GT)^{13}$ | 202 | F: GGTTAGTTAGCATATTCCCATCC-3’<br>R: TATCATCACAGCAATTACTCCTTC | |
| no Q solution | sarms0830 | VIC | $(AC)^{8}$ | 166 | F: AAGCAGAAATCAACACCTCTGTC<br>R: TAATGTGGCCTTGGATTCAGAAG | |
| | sarms3213 | VIC | $(CTAC)^{2}$ CAT $(CTAC)^{14}$ | 404 | F: CATTCTGGTCCCTGTGG<br>R: AATCATCTACCTCTTTACTG | |
| | sarms4545 | NED | $(ATT)^{2}$ $(GTT)^{7}$ | 238 | F: ACCACCACCATTCCCTTCTCCT<br>R: CATGCCTGGCCTTGCTTAG | |
| | sarms4547 | 6-FAM | $(AGAT)^{12}$ | 402 | F: CATCTCTAGACTTGCTGCCAATC<br>R: AGGAGCAGCAATATACCTCTCAG | |
| | sarms5711 | PET | $(AGAT)^{11}$ | 382 | F: TTACTGAAATTCCTCTGGTCCAG<br>R: TACTGAAATCTCTTGTGTGCAGC | |
| | sarms0445 | 6-FAM | $(CT)^{10}$ | 145 | F: AACAAGTCATGCCAGACTAAACC<br>R: AGAATATCATGGGGAACCTTGTC | Monomorphic |
| Multiplex 5 | sa52.20 | VIC | $(TTG)^{9}$ | 103-112 | From Chan et al. 2007 | |
| 60°C annealing | sa52.29 | NED | $(TTTC)^{15}$ | 185-249 | From Chan et al. 2007 | |
| no Q solution | sa60.02 | NED | $(GT)^{10}$ | 117-133 | From Chan et al. 2007 | |
| | sar80 | PET | $(ATAG)^{12}$ $(ACAG)^{6}$ | 349-402 | From Chan et al. 2007 | |
| | sar84 | PET | $(CA)^{16}$ | 109-147 | From Chan et al. 2007 | |

Supp. Mat., Table 2. Substitution models for EBSP analyses

| Phylogroup | mtDNA | PRLR | R35 | scar298 | scar875 |
|---|---|---|---|---|---|
| A | HKY | F81 | HKY+I | HKY+I | HKY |
| B | GTR | F81 | HKY | HKY+I | HKY |
| C | GTR | F81 | HKY | HKY+I | HKY |
| D | GTR+I | F81+I | HKY+I | GTR+I | F81+I |
| E | GTR+I | HKY | HKY+I | GTR+I | F81+I |

Supp. Mat., Table 3. Summary statistics for DNA sequence data.

| Locus | N | Alignment Length | Parsimony Informative Sites | Unique Seqs | Segregating sites, S | Average nucleotide differences, k | Nucleotide diversity, π |
|---|---|---|---|---|---|---|---|
| mtDNA | 225 | 2097 | 72 | 78 | 106 | 8.72 | 0.0043 |
| PRLR | 208 | 541 | 9 | 12 | 9 | 0.95 | 0.0018 |
| R35 | 195 | 658 | 5 | 12 | 8 | 0.71 | 0.0019 |
| scar298 | 213 | 341 | 13 | 14 | 13 | 1.92 | 0.0057 |
| scar875 | 212 | 287 | 12 | 20 | 13 | 1.27 | 0.0044 |

Supp. Mat., Table 4. Summary of microsatellite data for individuals from each of the eleven geographic regions.

| Region | N | Expected Heterozygosity | Observed Heterozygosity | Average gene diversity | Mean number of alleles | G-W Index | Mean Allelic richness |
|---|---|---|---|---|---|---|---|
| AA | 22 | 0.68755 | 0.63154 | 0.529 | 6.333 | 0.31496 | 2.393 |
| AB | 10 | 0.59348 | 0.51338 | 0.454 | 4.407 | 0.29101 | 2.171 |
| BA | 42 | 0.64902 | 0.57529 | 0.602 | 8.296 | 0.3371 | 2.560 |
| BB | 6 | 0.63409 | 0.41667 | 0.456 | 3.074 | 0.26077 | 2.103 |
| C | 20 | 0.65211 | 0.61647 | 0.647 | 7.593 | 0.31783 | 2.624 |
| DA | 76 | 0.6962 | 0.64657 | 0.684 | 10.815 | 0.37527 | 2.707 |
| DB | 16 | 0.64172 | 0.63093 | 0.593 | 6.667 | 0.3398 | 2.579 |
| EA | 27 | 0.67652 | 0.60591 | 0.649 | 7.852 | 0.34462 | 2.592 |
| EB | 10 | 0.66979 | 0.61111 | 0.670 | 5.667 | 0.33515 | 2.622 |
| EC | 8 | 0.73487 | 0.64263 | 0.653 | 5.667 | 0.32679 | 2.770 |

**Supplementary Material Figure Legends**

Supp. Mat., Figure 1. Alternative population models evaluated with approximate Bayesian computation methods.

Supp. Mat., Figure 2. Haplotype networks for each of the nuclear genes sequenced. Circles represent unique alleles with the size of the circle corresponding to the relative abundance and the color referring to the region of origin of individuals with that haplotype. Lines connecting haplotypes represent one mutational step. Small white circles represent unsampled haplotypes. [Alternate version for individuals with color vision deficiencies included].

Supp. Mat., Figure 3. Extended Bayesian Skyline plots based on mtDNA and nuclear sequence data for individuals from regions A, B, C, D, and E.

### Supplementary Materials, Figure 1

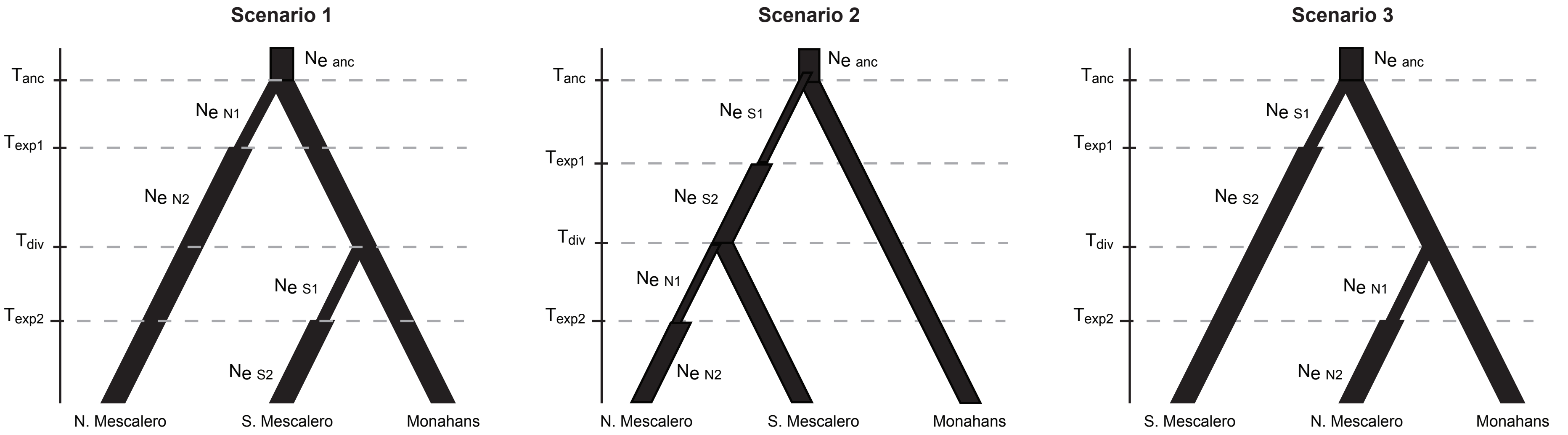

**Supplementary Materials, Figure 2**

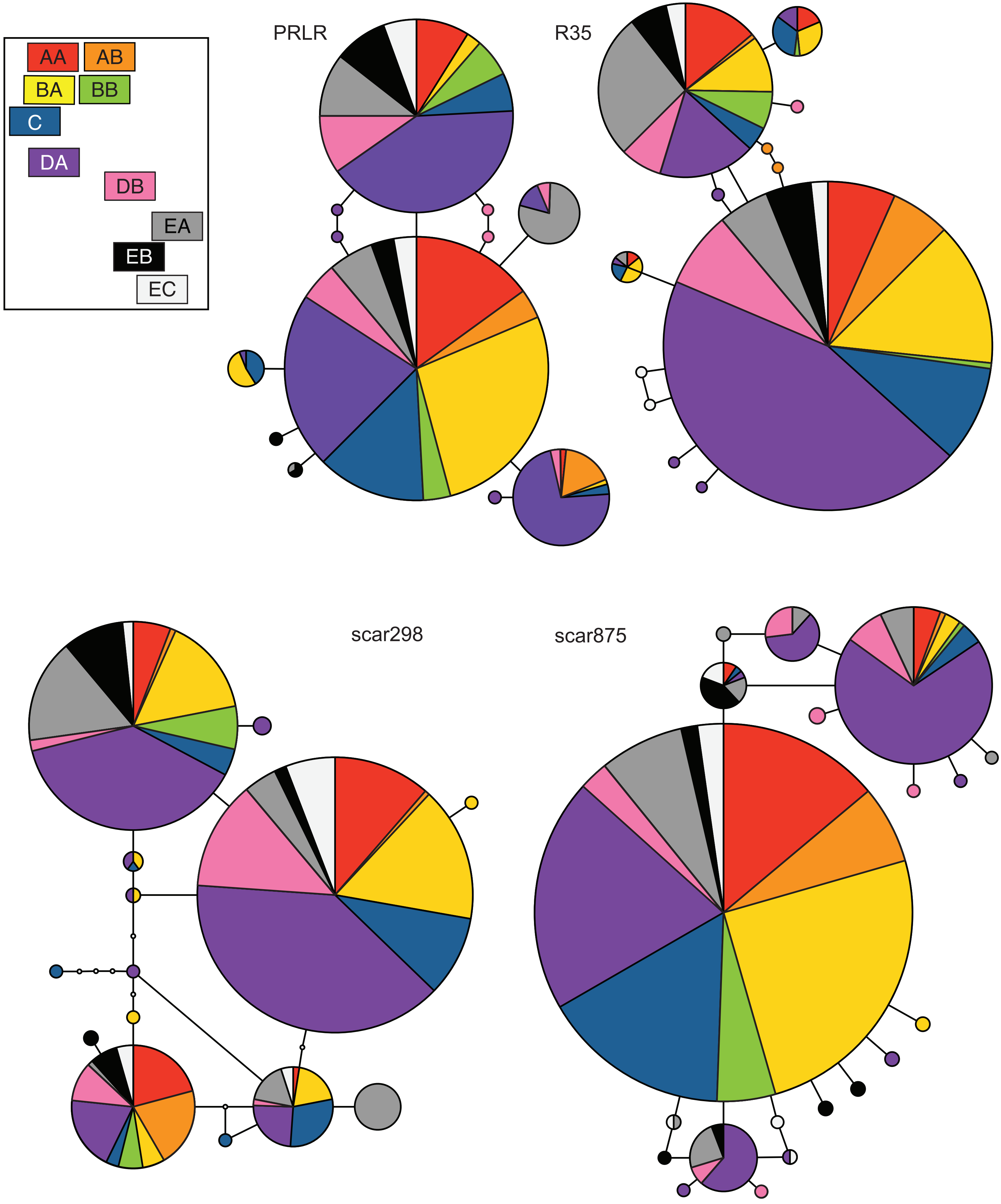

**Supplementary Materials, Figure 2 (alternate version)**

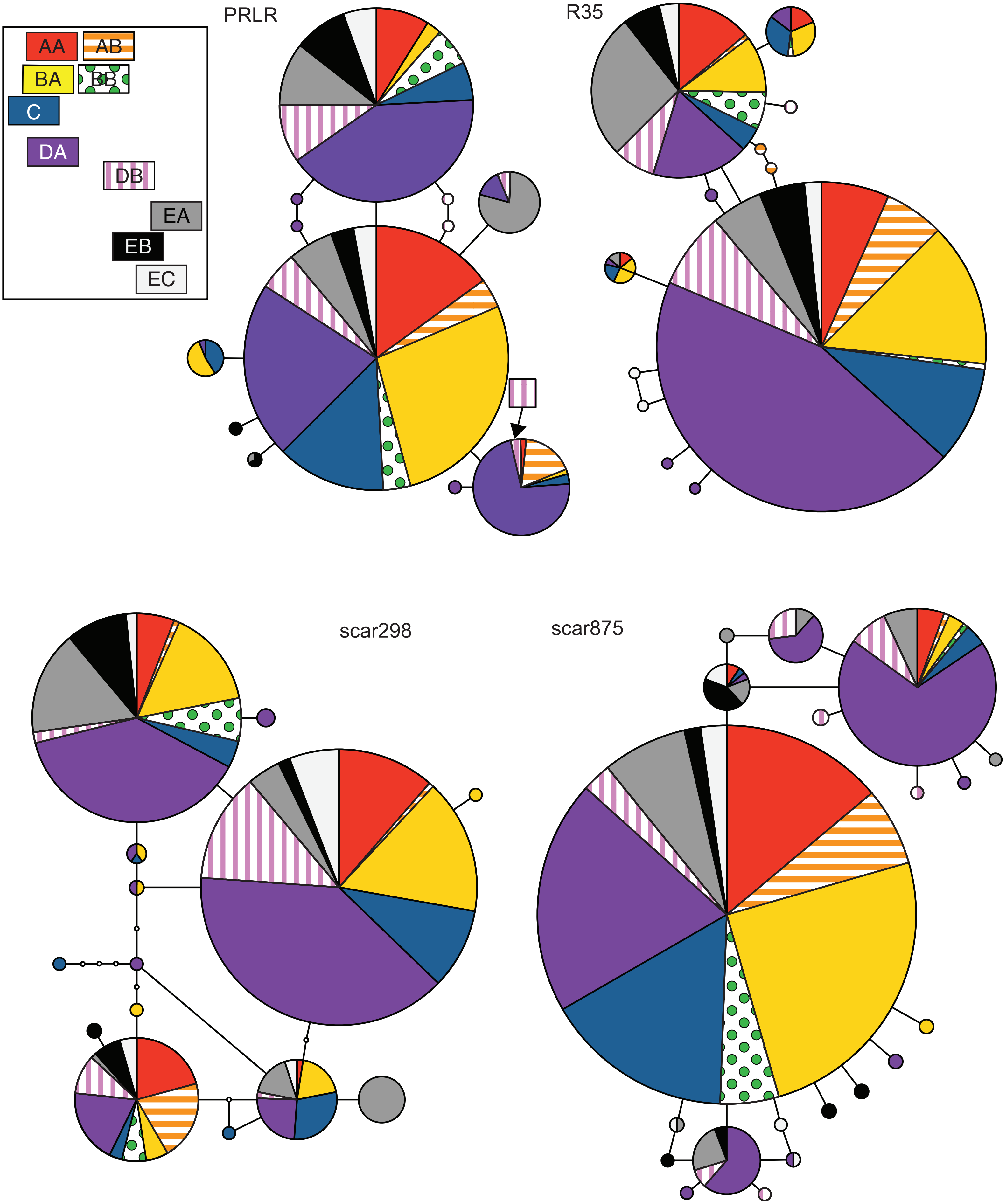

#### Supplementary Materials, Figure 3

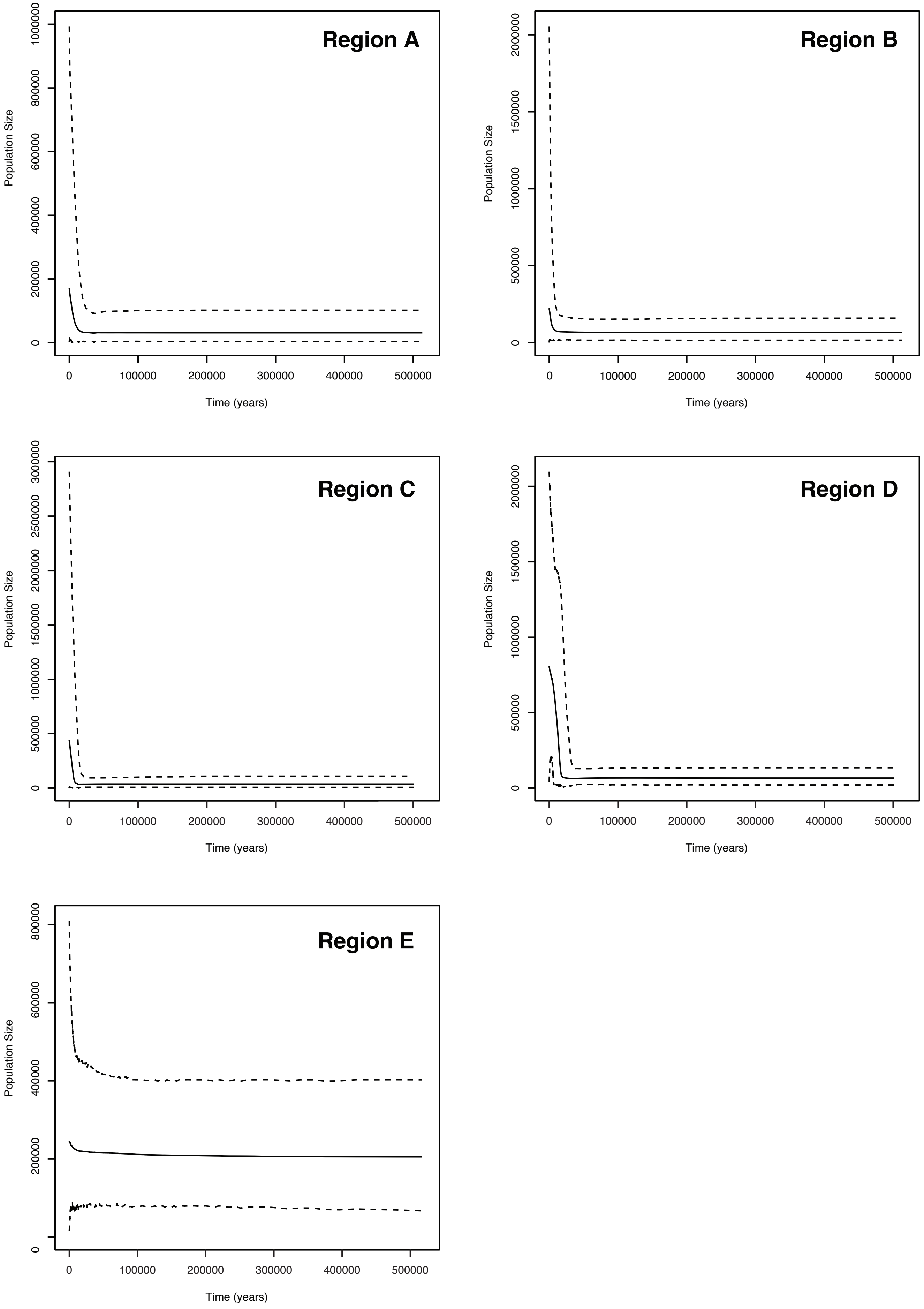
